## Supplementary Figures for "Social isolation modulates appetite and defensive behavior via a common oxytocinergic circuit in larval zebrafish"

### SUPPLEMENTARY FIGURES & LEGENDS:

### SUPPLEMENTARY FIGURE 1

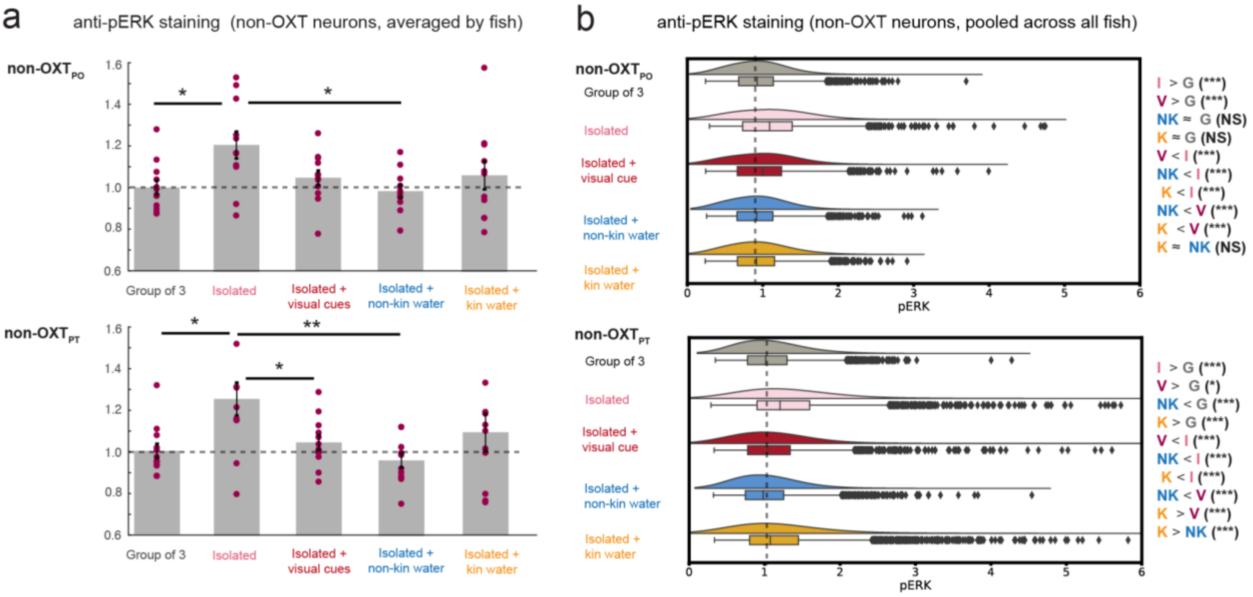


**Supplementary Figure 1: Modulation of nearby non-OXT neurons by social context**

**(a)** Effect of social isolation and social sensory cues on mean non-OXT$_{PO}$ and non-OXT$_{PT}$ neuron activity per fish. pERK intensities shown are normalized to the mean PO or PT intensities in control fish (i.e. fish maintained in groups of 3). Social isolation significantly increased the activity of non-OXT neurons in the PO and PT, and visual, non-kin and kin cues were sufficient to restore OXT activity, with non-kin water showing the most significant effect on both non-OXT$_{PO}$ and non-OXT$_{PT}$ neurons.

**non-OXT$_{PO}$ neurons:** p = 0.034* (group vs isolated) / 0.29 (group vs visual) / 0.93 (group vs non-kin water) / 0.74 (group vs kin water) / 0.131 (isolated vs visual) / 0.026* (isolated vs non-kin) / 0.066 (isolated vs kin water) / 0.207 (visual vs non-kin water) / 0.34 (visual vs kin water) / 0.74 (kin vs. non-kin water).

**non-OXT$_{PT}$ neurons:** p = 0.015* (group vs isolated) / 0.26 (group vs visual) / 0.64 (group vs non-kin water) / 0.34 (group vs kin water) / 0.024* (isolated vs visual) / 0.0039** (isolated vs non-kin) / 0.13 (isolated vs kin water) / 0.10 (visual vs non-kin water) / 0.78 (visual vs kin water) / 0.19 (kin vs. non-kin water).

**(b)** Probability distributions and box plots showing median, inter-quartile range and 95% confidence interval of normalized pERK fluorescence across all neurons per group. pERK fluorescence was divided by the mean pERK fluorescence of all non-OXT neurons (PO and PT combined) of the control group. Fish were either kept in groups of 3 (gray, n = 6700 non-OXT$_{PO}$ / 5678 non-OXT$_{PT}$ neurons from 12 fish), isolated (pink, n = 6594 non-OXT$_{PO}$ / 4962 non-OXT$_{PT}$ neurons from 11 fish), isolated but exposed to visual cues of conspecifics through a transparent barrier (red, n = 5157 non-OXT$_{PO}$ / 4081 non-OXT$_{PT}$ neurons from 12 fish) or isolated but exposed to non-kin-conditioned water (blue, n =5099 OXT$_{PO}$ / 4235 OXT$_{PT}$ neurons from 11 fish) or kin-conditioned water (orange, n= 4574 non-OXT$_{PO}$ / 4621 non-OXT$_{PT}$ neurons from 12 fish).

**OXT$_{PO}$ neurons:** p = $8.4 \times 10^{-107}$*** ( group vs isolated) / $7.2 \times 10^{-17}$*** (group vs visual) / 0.071 (group vs non-kin water) / 0.98 (group vs kin water) / $9.2 \times 10^{-36}$*** (isolated vs visual) / $1.2 \times 10^{-106}$*** (isolated vs non-kin water) / 6.6x10$^{-8}$*** (isolated vs kin water) / 9.5x10$^{-21}$***(visual vs non-kin water) / 7.8x10$^{-13}$*** (visual vs kin water) / 0.11 (kin vs non-kin water). Two-sided Wilcoxon Rank-Sum test.

**OXT$_{PT}$ neurons:** p = 2.9x10$^{-109}$*** ( group vs isolated) / 0.022* (group vs visual) / 2.3x10$^{-5}$*** (group vs non-kin water) / 7.9x10$^{-19}$*** (group vs kin water) / 4.4x10$^{-70}$*** (isolated vs visual) / 1.2x10$^{-129}$*** (isolated vs non-kin water) / 3.2x10$^{-30}$*** (isolated vs kin water) / 4.0 x 10$^{-9}$*** (visual vs non-kin water) / 2.4x10$^{-9}$***(visual vs kin water) / 1.6x10$^{-32}$*** (kin vs non-kin water). Two-sided Wilcoxon Rank-Sum test.

### SUPPLEMENTARY FIGURE 2

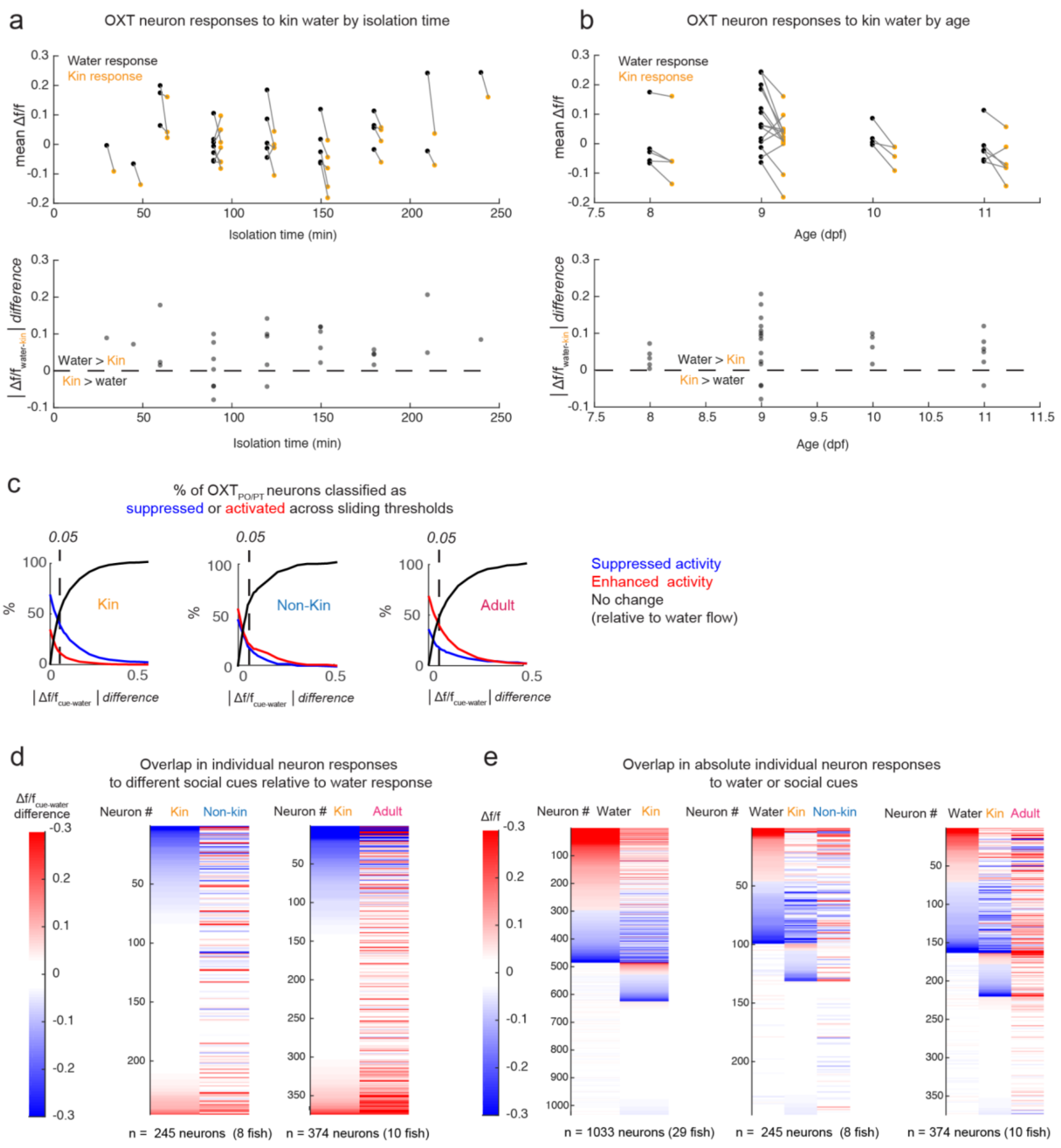


**Supplementary Figure 2: Additional characterization of OXT neuron responses to social cues and water flow**

a) Effect of kin water on OXT neuron activity plotted as a function of isolation time (min). **Top panel:** Average post-stimulus calcium activity per fish (n = 29 fish) in response to either water flow (black) or kin water (orange). **Bottom panel:** Difference in water and kin water-induced calcium activity per fish. Fish were isolated for between 30 min to 4 hrs before imaging, though the bulk of experiments were performed with 1.5 - 2.5 hr isolation (note that time of isolation = time of embedding in agarose).

b) Effect of kin water on OXT neuron activity plotted as a function of age (dpf). **Top panel:** Average post-stimulus calcium activity per fish (n = 29 fish) in response to either water flow (black) or kin water (orange). **Bottom panel:** Difference in water and kin water-induced calcium activity per fish.

c) Percentage of OXT neurons that would be classified as suppressed (blue) or activated (red) by each water-borne cue, as a function of the mean difference in integrated calcium activity from the water response (i.e., difference threshold). We quantified each neuron's activity as the difference in integrated Δf/f within a 60 second window surrounding the stimulus (post minus pre). For each social cue, we then subtracted the same neuron's average response to the control stimulus (i.e., water flow) in order to normalize each neuron's social cue response *relative to water flow*. A difference threshold of 0.05 (i.e., 5% difference) was used in subsequent panels.

d) OXT neurons sorted according to their differential response to kin water (relative to water flow) and their corresponding activities in response to non-kin or adult water. The data here is similar to **Figure 2e**, but not categorized into "suppressed" or "activated" classes.

e) Absolute calcium response (post minus pre-stimulus integrated Δf/f) of OXT neurons to water flow and other social cues, sorted according to their response to water flow.

### SUPPLEMENTARY FIGURE 3

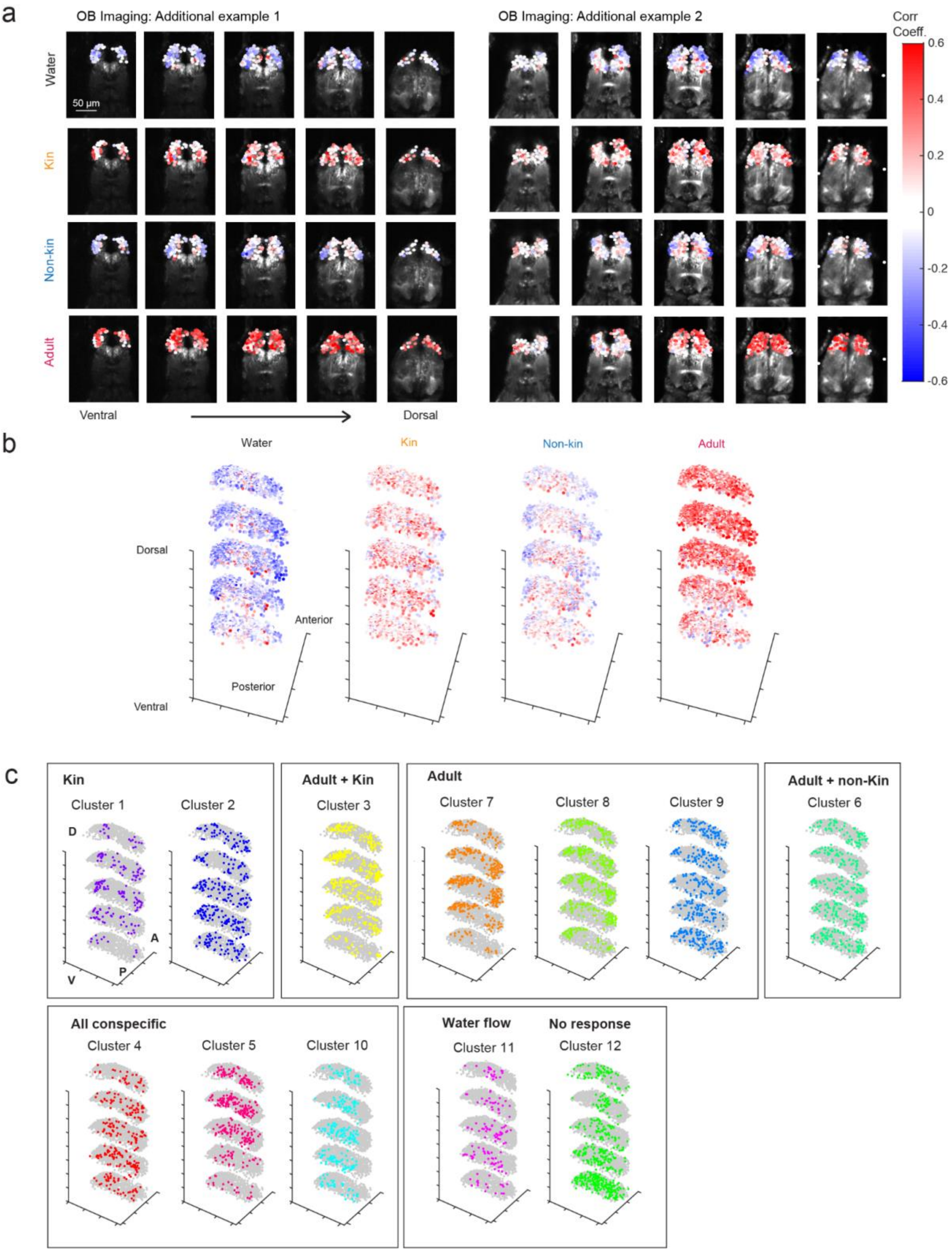


**Supplementary Figure 3: The zebrafish olfactory bulb discriminates conspecific cues**

**(a)** Units extracted from two additional fish color coded (blue to red) by their coefficient correlations to each stimulus, and overlaid over anatomy images, per z-plane. Scale bar = 50 µm.

**(b)** 3D plot displaying correlation coefficients of all units across all fish to each stimulus. Color code same as in (a). As in Figure 3, x-y coordinates for all units per fish was scaled linearly to their minimum and maximum values in each dimension.

**(c)** 3D plot displaying the spatial localization of units within each cluster. A = Anterior, P = Posterior, D = Dorsal, V = Ventral.

### SUPPLEMENTARY FIGURE 4

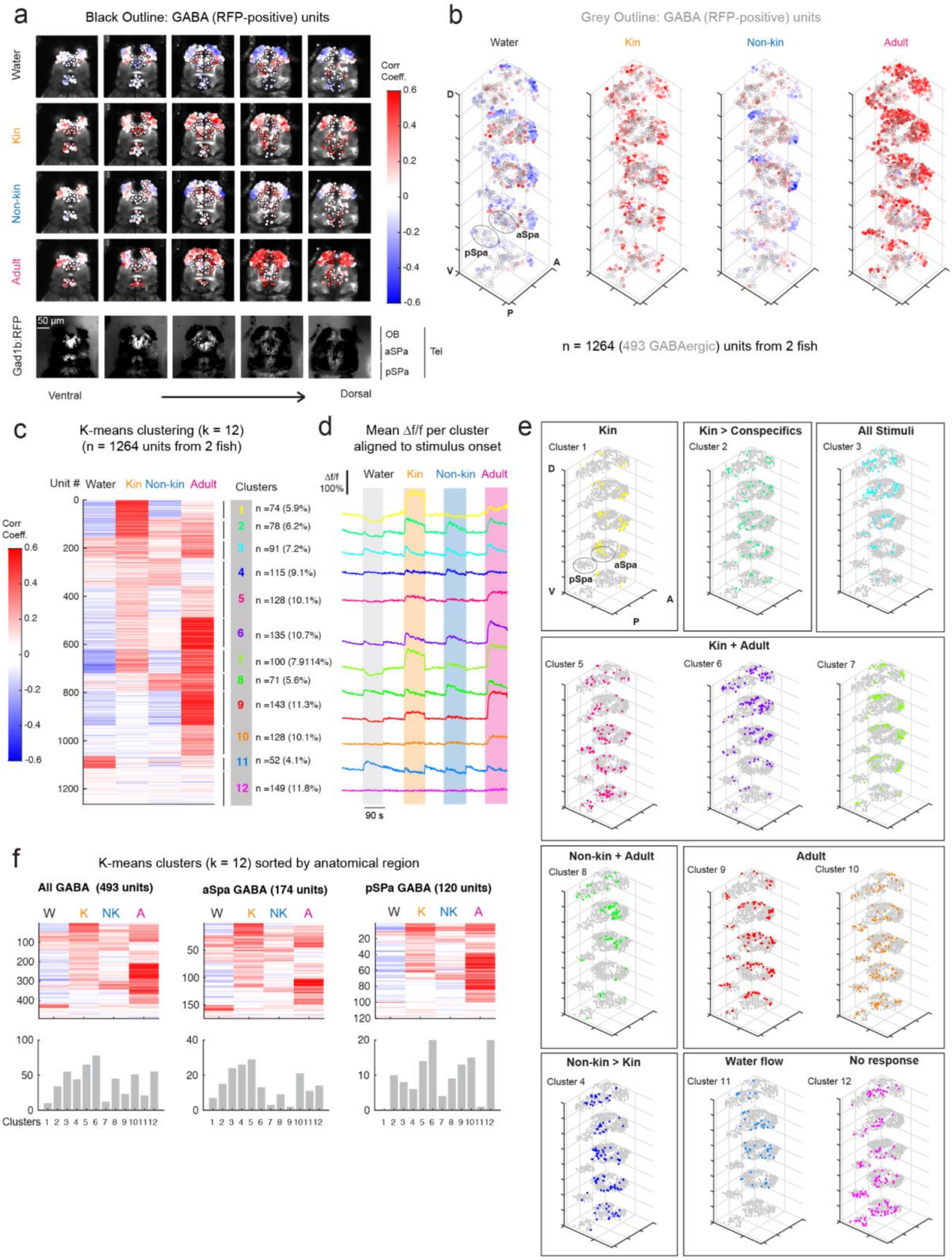


**Supplementary Figure 4: GABAergic subpallial neurons respond to conspecific stimuli**

a) Calcium imaging was performed on the telencephalon (including both OB and SPa) of transgenic fish (n = 2) co-expressing *Tg(HuC:GCaMP6s)* and *Tg(Gad1b:RFP)* as they were exposed to water flow, kin, non-kin, or adult water. Automatedly segmented units were identified as GABAergic (black outline) based on thresholded red fluorescence intensities (see **Methods**). Correlation coefficients for each unit with each stimulus regressor have been color coded (blue to red) and overlaid over the anatomy image per z-plane. Bottom-most row shows an anatomy stack of GABAergic cells per z-plane. GABAergic cells are present in the olfactory bulb, as well as anterior and posterior subpallium, which is part of the ventral telencephalon. Scale bar = 50 µm.

b) 3D plot displaying correlation coefficients of all units from all fish to each stimulus. Color code is the same as in (a). As in Figure 2, XY coordinates for all units per fish were scaled linearly to their minimum and maximum values in each dimension. GABAergic units are outlined in grey.

c) Correlation coefficients of all neurons (1264 units from 2 fish) to each stimulus, sorted according to K-means cluster (k = 12). Number of units within each cluster and percentage representation of total units are displayed on the right.

d) Mean stimulus-triggered activity (Δf/f) for each cluster, aligned to stimulus onset. Order of clusters and units are the same as in (c).

e) 3D plot displaying the spatial localization of units within each cluster. Color code same as in (d).

f) After k-means clustering (as in (c)), units were segmented into putative anatomical regions based on GABAergic identity and/or anterior-posterior location (aSPa located anteriorly but posterior to the OB, and pSPa cluster is posterior to the anterior commissure). Bar graphs show the number of units within each anatomical region belonging to each cluster.

### SUPPLEMENTARY FIGURE 5

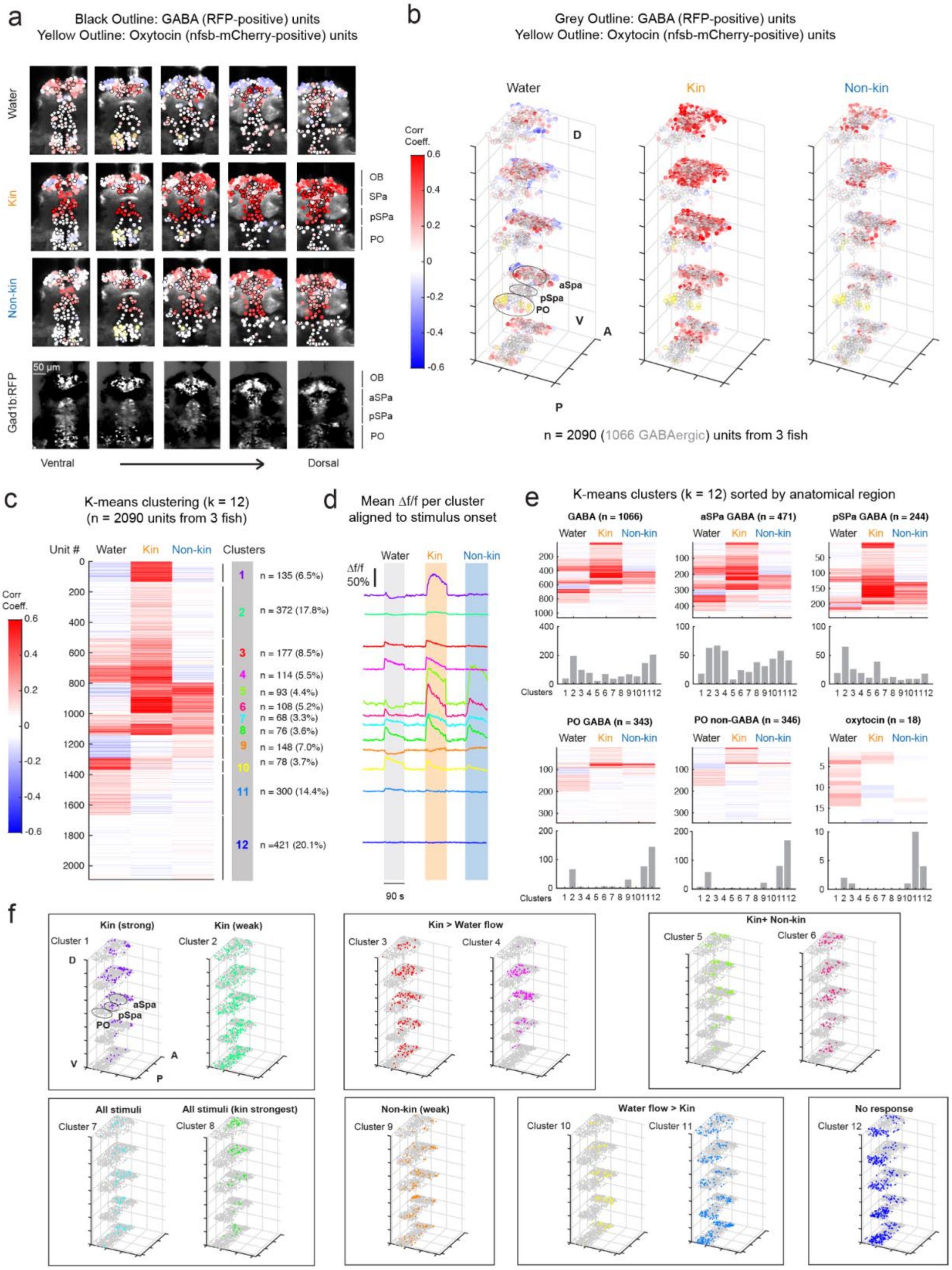


**Supplementary Figure 5: GABAergic subpallial neurons discriminates kin and non-kin cues**

a) Calcium imaging was performed on the telencephalons of transgenic fish (n = 3) co-expressing *Tg(HuC:GCaMP6s), Tg(Gad1b:RFP), and Tg(oxt:Gal4; UAS:nfsb-mCherry)*, as they were exposed to water flow, kin or non-kin water. Segmented units were identified as GABAergic (black outline) based on thresholded red fluorescence, whereas putative oxytocin neurons (yellow outline) were manually identified based on their brighter, more punctated labeling. Correlation coefficients for each unit with each stimulus regressor are color-coded (blue to red) and overlaid over the anatomy image per z-plane. Bottom-most row shows an anatomy stack of GABAergic cells per z-plane. GABAergic cells are present in the olfactory bulb, subpallium, and preoptic area. Scale bar = 50 µm.

b) 3D plot displaying correlation coefficients of all units across all fish to each stimulus. Color code same as in (a). XY coordinates for all units per fish were scaled linearly to their minimum and maximum values in each dimension. GABAergic units outlined in grey.

c) Correlation coefficients of all neurons (2090 units from 3 fish) to each stimulus, sorted according to K-means cluster (k = 12). Number of units within each cluster and percentage representation of total units are displayed on the right. Clusters specific to kin water and adult water, as well as clusters with mixed selectivity were observed. Clusters 1-4 show stronger responsiveness to kin cues over other cues. Note that clusters 5-8 are dominated by a single fish (shown in (a)) that had slightly stronger non-kin responses, the other 2 fish had weak non-kin responses (e.g. cluster 9).

d) Mean stimulus-triggered activity (Δf/f) for each cluster, aligned to stimulus onset. Order of clusters and units are the same as in (c).

e) After k-means clustering (as in (c)), units were segmented into putative anatomical regions based on GABAergic identity and/or anterior-posterior location (aSPa located anteriorly but posterior to the OB, and pSPa cluster is posterior to the anterior commissure). Bar graphs show the number of units within each anatomical region belonging to each cluster. Note that the PO region (including OXT neurons), unlike the SPa, predominantly has a stronger water flow response relative to kin or non-kin cues.

f) 3D plot displaying the spatial localization of units within each cluster. Color code same as in (e).

### SUPPLEMENTARY FIGURE 6

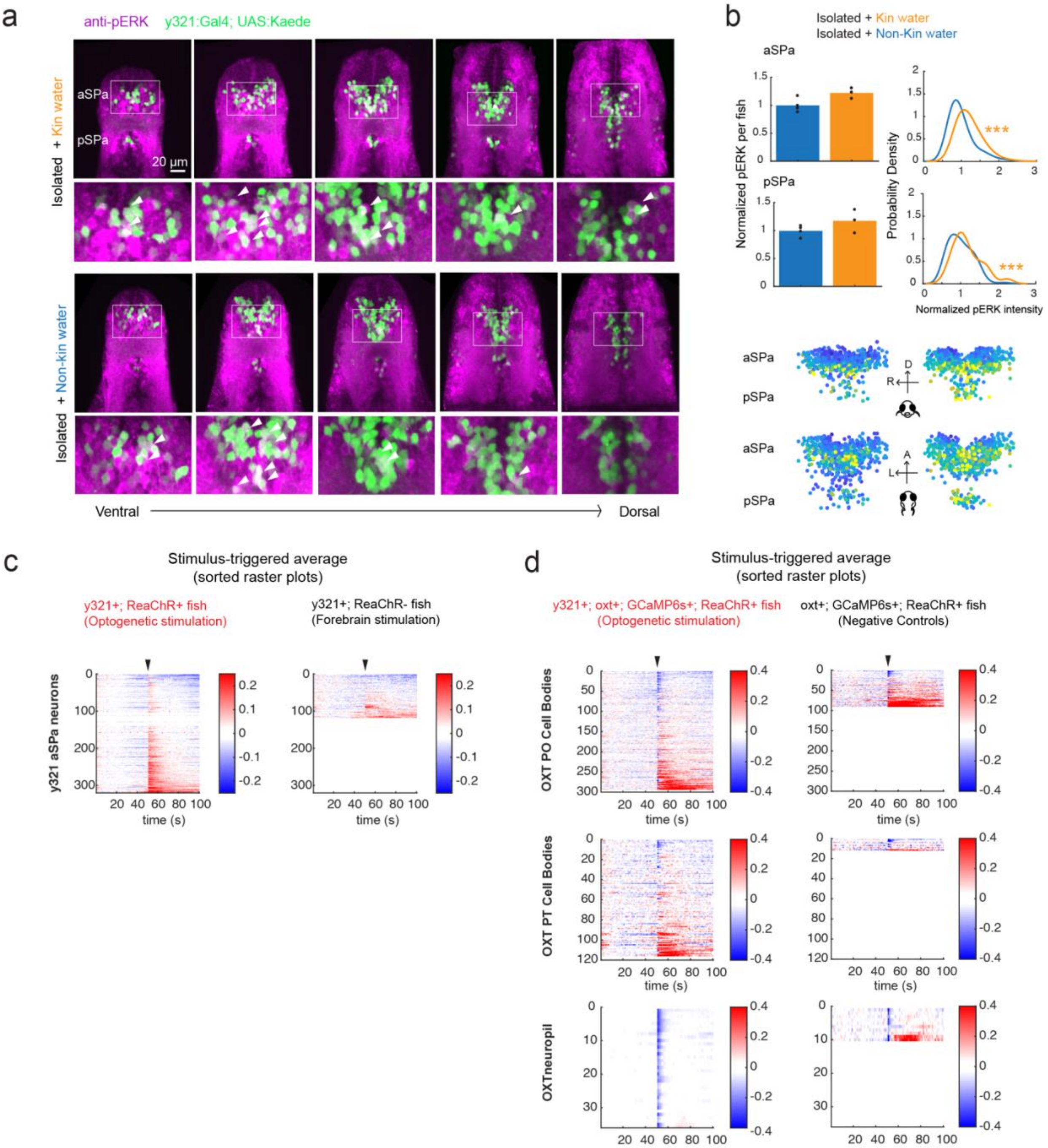


**Supplementary Figure 6: y321 subpallial neurons are responsive to conspecific cues and sufficient to suppress OXT neuron activity**

(a) A subset of subpallial y321 cells overlap with anti-pERK staining following exposure to kin **(top)** or non-kin **(bottom)** water. Insets show the aSPa y321 neurons at higher magnification. Scale bar = 20 µm.

(b) **Top:** Quantification of pERK fluorescence in aSPa and pSPa y321 neurons after exposure to kin and non-kin water. When comparing the mean pERK signal per fish, there was no significant difference between kin (n= 3 fish) and non-kin (n = 4 fish) responses (aSPa: p = 0.11; pSPa: p = 0.40, Two-sided Wilcoxon Rank-Sum Test). However, there was a significant right shift in pERK intensities in response to kin versus non-kin odors (aSPa: ***p = $6.2x10^{-42}$, n 967 neurons (non-kin) / 637 neurons (kin)); pSPa: ***p = $1.4x10^{-4}$, n = 100 neurons (non-kin), n = 72 neurons (kin), Two-sided Wilcoxon Rank-Sum Test). **Bottom:** Spatial distribution of y321 SPa neurons. Both the front view and top view are depicted (fish orientation shown in schematic), color coded by pERK intensity (yellow = highest intensity blue = lowest intensity). 200 y321 neurons were randomly sampled per fish, with 600 neurons were drawn from this sample, to ensure even representation across all fish and groups.

(c) Raster plots showing responses of individual y321 positive neurons to y321 optogenetic stimulation (**left**) or a control stimulation (**right;** when ReaChR is not expressed). Laser pulse occurs at the 50 s mark (indicated by black arrow).

(d) Raster plots showing responses of individual OXT neurons in the PO or PT, or OXT neuropil regions, to y321 optogenetic stimulation (**left**) or a control stimulation (**right;** when y321:Gal4 is not expressed). Laser pulse occurs at the 50 s mark (indicated by black arrow).

### SUPPLEMENTARY FIGURE 7

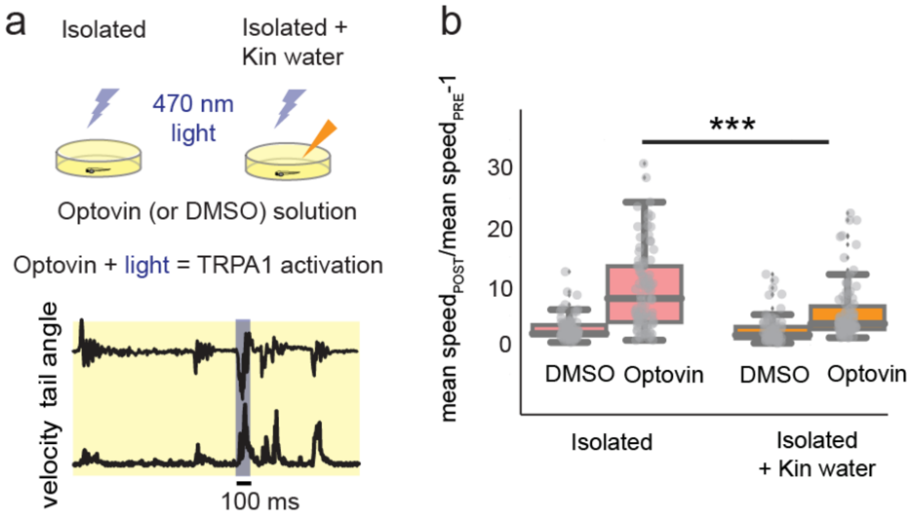


**Supplementary Figure 7: Social context affects defensive behavior in larval zebrafish**

**(a)** Schematic for how the effect of kin-conditioned water on nocifensive behavior was tested. Fish were incubated with DMSO or Optovin solution, with or without kin-conditioned water. Short illumination with a 470 nm LED activates TRPA1 receptors and elicits large-angle tail bends and a corresponding increase in swim velocity.

**(b)** The presence of conspecific cues significantly decreases TRPA1-induced velocity changes ***$p<0.001$, two-sided Wilcoxon Rank-Sum test.

### SUPPLEMENTARY FIGURE 8

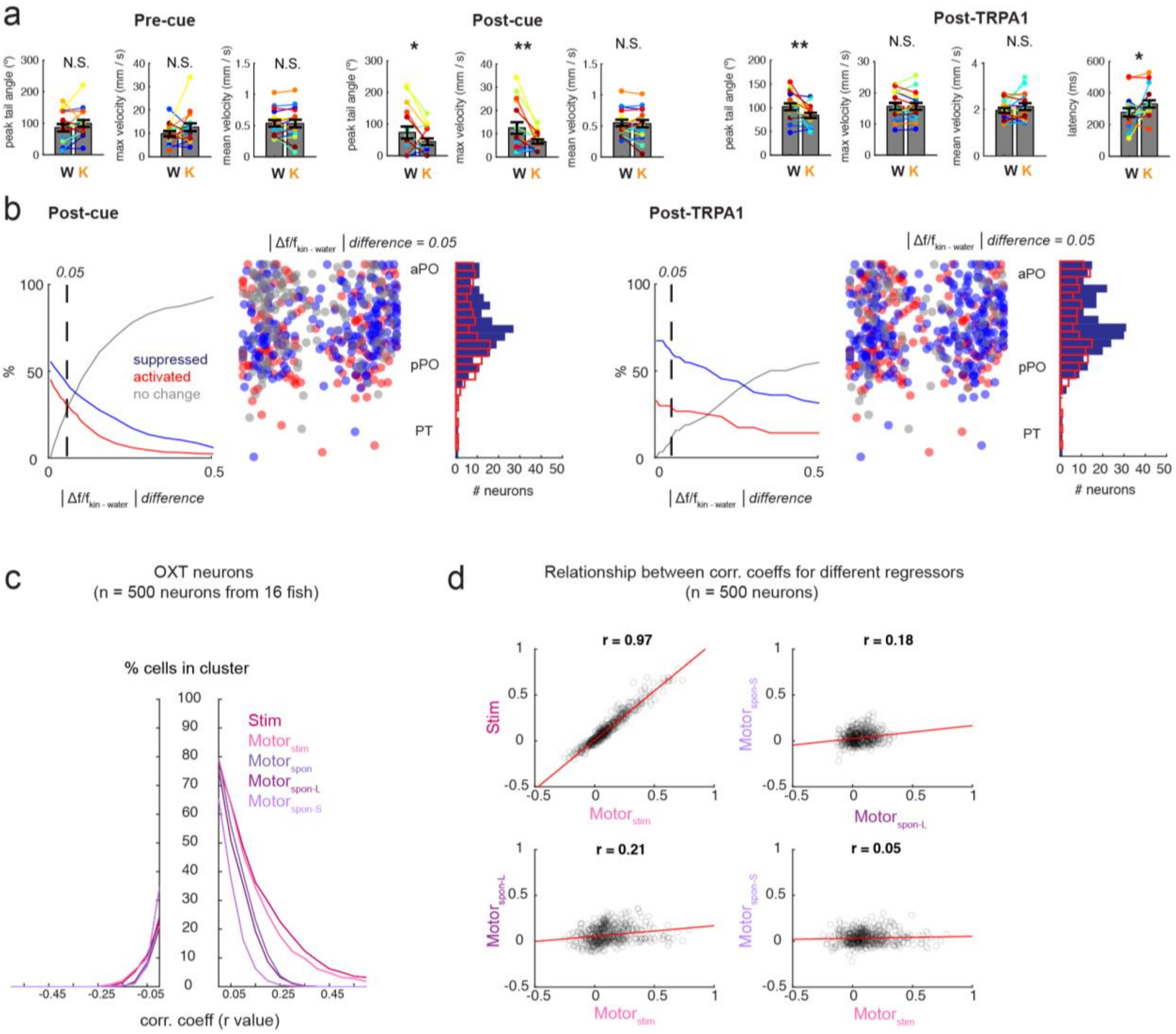


**Supplementary Figure 8: Effect of kin water on behavior and OXT neuron activity**

(a) Effect of kin water on kinematics of swim bouts that occur either prior to cue delivery, post-cue delivery or directly after TRPA1 stimulation. Peak tail angle: p = 0.95 / 0.038* / 0.0061**; Maximum velocity: p = 0.89 / 0.0075** / 0.51; Mean velocity: 0.67 / 0.51 / 0.80, response latency (post-TRPA1 stimulation): p = 0.047*, one-sided Wilcoxon Signed-Rank test.

(b) **Left:** Percentage of OXT neurons that would be classified as suppressed (blue) or activated (red) by kin water, either pre- or post-TRPA1 stimulation, as a function of the mean difference in integrated calcium activity from the water response (i.e., difference threshold). **Center:** spatial distribution of suppressed and activated neurons. **Right:** Distribution of suppressed and activated neurons along the A-P axis, using a threshold of 0.05 (5% difference).

(c) Percentage of cells which would be classified as being correlated with each regressor as a function of correlation coefficient, demonstrating strong OXT neuron correlation with TRPA1 stimulus and stimulus-induced behavior, and weaker correlation to spontaneous motor events.

(d) Relationship between each OXT neuron's correlation coefficients to different regressors.

### OTHER SUPPLEMENTARY LEGENDS:

**Supplementary Movie 1:** Z-stack (dorsal to ventral) of brain activity map shown in **Figure 1a**, overlaid on a *Tg(etVMAT:GFP)* brain used as an anatomical reference. Scale bar = 50 µm.

**Supplementary Data 1:** Z-brain anatomical regions that are more activated in isolated fish as compared to fish in a group. No regions were identified showing the opposite pattern (i.e., less active in isolated fish).
